# A simple, robust, broadly applicable insertion mutagenesis method to create random fluorescent protein – target protein fusions

**DOI:** 10.1101/2024.01.30.578020

**Authors:** Andrew Pike, Cassandra Pietryski, Padraig Deighan, Jason Kuehner, Derek Lau, Anupama Seshan, Paul E. March

## Abstract

A simple broadly applicable method was developed using an *in vitro* transposition reaction followed by transformation into *Escherichia coli* and screening plates for fluorescent colonies. The transposition reaction catalyzes the random insertion of a fluorescent protein open reading frame into a target gene on a plasmid. The transposition reaction is employed directly in an *E. coli* transformation with no further procedures. Plating at high colony density yields fluorescent colonies. Plasmids purified from fluorescent colonies contain random, in-frame fusion proteins into the target gene. The plate screen also results in expressed, stable proteins. A large library of chimeric proteins was produced that was useful for downstream research. The effect of using different fluorescent proteins was investigated as well as the dependence of linker sequence between the target and fluorescent protein open reading frames. The utility and simplicity of the method was demonstrated by the fact that it has been employed in an undergraduate biology laboratory class without failure over dozens of class sections. This suggests that the method will be useful in high impact research at small liberal arts colleges with limited resources. However, in frame fusion proteins were obtained from eight different targets suggesting that the method is broadly applicable in any research setting.

**Summary:** This report describes a simple screen to obtain random insertions of fluorescent proteins into a target protein of interest. The screen results in a useful library of mutated and functional tagged proteins that can be employed to investigate protein biochemical activity, protein structure, protein function and protein cellular localization. The screen is robust, generally applicable, and has been employed in an undergraduate laboratory class. It is also useful for studies in advanced research-intensive projects carried out in institutions of all types.

**Graphical Abstract:** 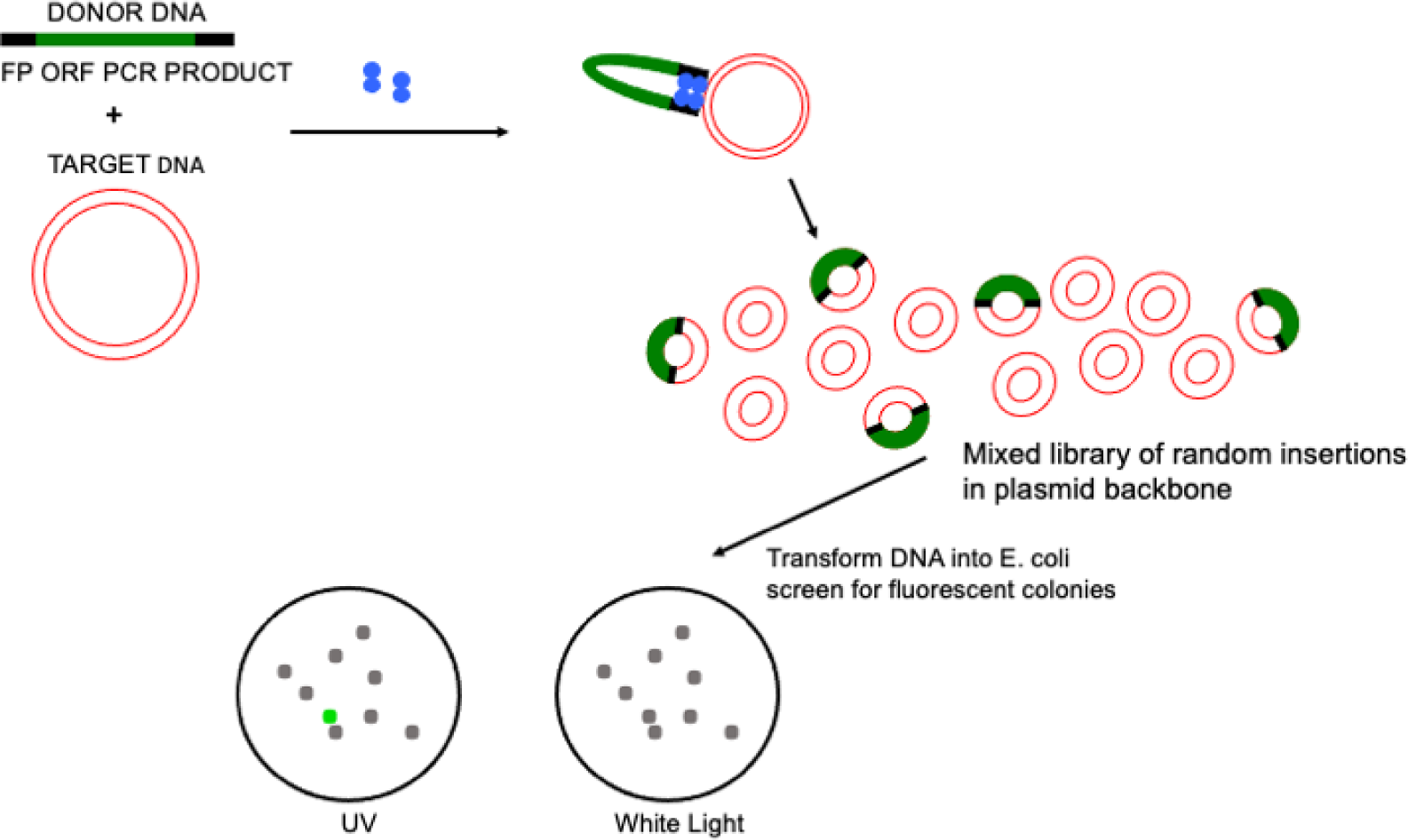

## Introduction

High impact contemporary research in molecular biology and molecular genetics has traditionally been carried out at institutions with significant budgets for research in labs staffed with graduate students and postdoctoral researchers. Opportunities for undergraduate participation were mostly limited to individual opportunities to apprentice in research labs. Starting in the early 2000’s widespread efforts to remove barriers to research by undergraduates have been initiated. Research was shown to increase the depth of learning and a feeling of inclusion by students (1). There are still barriers to research by undergraduates. High impact research in some areas (including molecular biology and molecular genetics) often requires expensive reagents and equipment that may be beyond the reach of some institutions, especially small liberal arts colleges. A method that is simple, inexpensive, with broad application would help to remove barriers to research. Importantly, a wide-ranging approach should be useful in high impact research carried out at any level of research at any institution.

Here we describe an insertional mutagenesis method, dubbed MORFIN (mutagenesis by open reading frame insertion), that gives rise to fully functional mutants tagged with a fluorescent marker. The approach creates open reading frame (ORF) fusions between a fluorescent protein (FP) and a target protein. The method requires one *in vitro* step, and it results in a library of in-frame fusion proteins. Following transformation of this library into *E. coli,* it is possible to screen directly for fluorescent colonies with functional ORF fusions.

## Materials and Methods

The method described here depends on the activity of EZ-Tn5 Transposase (EZ-Tn5 is commonly sold in kits for specific purposes. The enzyme by itself, not part of a kit can be purchased from LGC Biosearch Technologies, Petaluma, CA). This enzyme only requires a 19-nucleotide inverted repeat at each end of a linear blunt-end DNA fragment to catalyze the random insertion of the DNA fragment into a target DNA. PCR Primers were designed to create amplicons that contained the 19-nucleotide inverted repeats at the 5’ and 3’ end of the amplicons (Table1). A FP ORF was included between the inverted repeats. Primer design included alteration of the FP’s ORF start codon and stop codon such that a single continuous ORF was present from the first to the last nucleotide on the amplicon. The details of these DNA primer modifications are shown in Table 1. Any DNA polymerase that results in blunt end amplicons can be employed in PCR reactions, in this work Phusion polymerase (Thermo Fisher Scientific) was employed. In order to increase robustness of PCR reactions using primers with longer ‘tails’ an initial PCR reaction using TAQ DNA polymerase was employed. Then 1 ul of the initial PCR reaction was employed in a second PCR using Phusion polymerase to create high-quality blunt-end amplicons. Amplicons from PCR reactions were purified using QIAquick PCR purification kit.

**Table 1:**
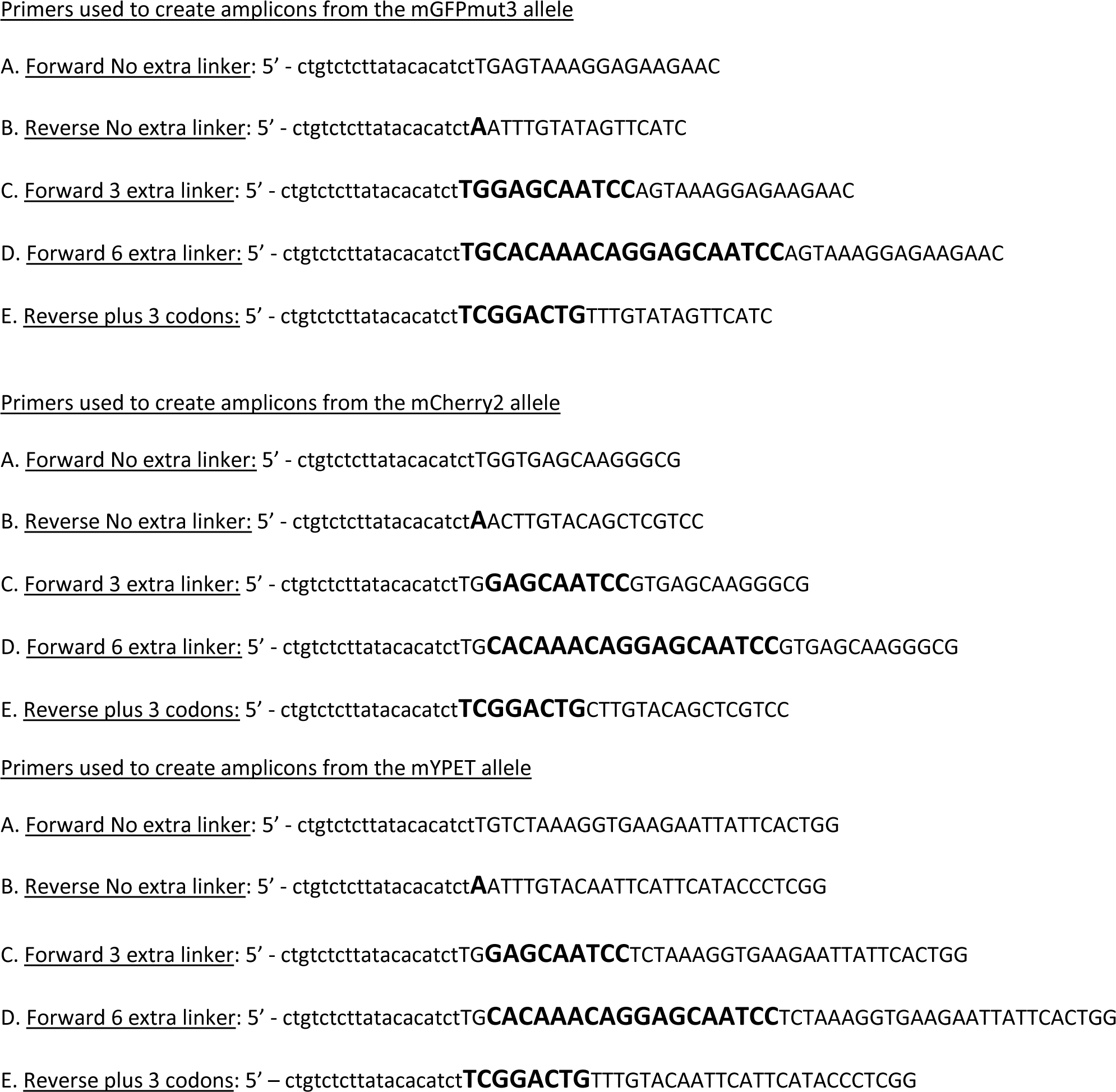
PCR primers employed to produce FP ORF amplicons with additional 5’ and 3’ flanking sequence. Lower case text denotes the 19-nucleotide Tn5 recognition element. Upper case black letters denote DNA sequence derived from the FP ORF. Upper case bold letters denote nucleotides that were altered to adjust the reading frame and inserted to add linker codons. The meaning of ‘no extra linker’, ‘3 extra linker’, and ‘6 extra linker’ is described in Figure 4. To create FP amplicons with no extra linkers, primer A and B were employed. To create an amplicon with 3 extra linkers primer C and B were employed. To create an amplicon with 6 extra linkers primer D and E were employed.

Target ORFs were contained on *E. coli* expression vectors (Table 2). Purified plasmid DNA was prepared using QIAprep Spin Miniprep kits. According to EZ-Tn5 manufacturer’s specifications it is essential to use high quality DNA (both amplicon and plasmid) for the transposition to work. The purity and concentration of DNA samples must be accurately determined. It is important to rigorously follow manufacturer’s instructions for the transposition reaction. Two hundred nanograms of target plasmid must be used and a molar ratio of amplicon – target DNA of 1:1 is required to ensure that multiple amplicons are not inserted into the same plasmid. The transposition reaction volume was always 10 ul. The reactions were incubated for 2 h at 37°C. Reactions were terminated using 1 ul of Stop solution supplied by the manufacturer and incubation at 70 °C for 10 min. Reactions were cooled on ice and 1 ul of this reaction mixture was used directly to transform *E. coli* MAX Efficiency DH5alpha competent cells (Thermo Fisher). Figure 1 shows an overview of the method and File S1 provides a step-by-step protocol.

**Table 2:**
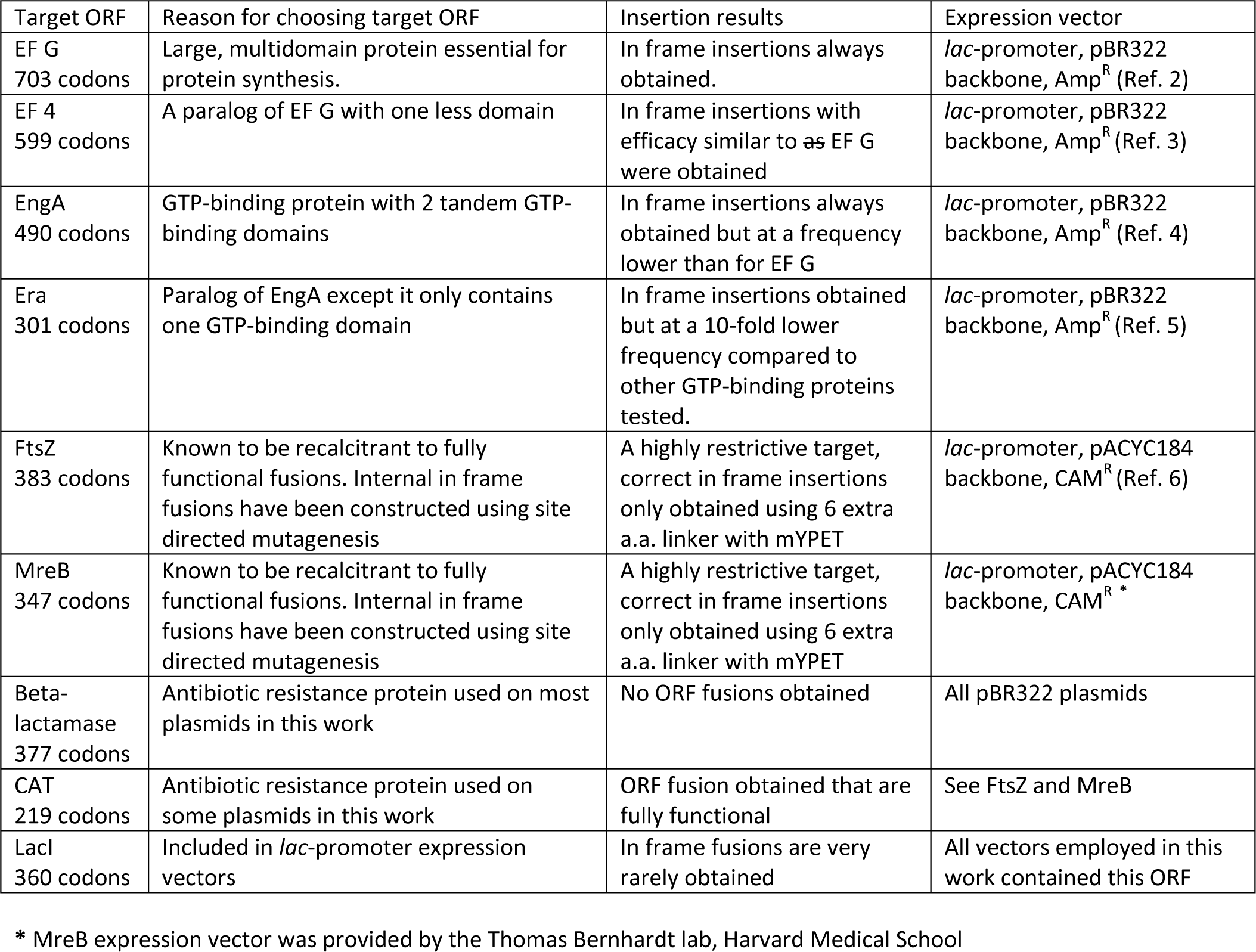
Summary of Target Protein ORFs. Transposition reactions were employed in transformations into ultra-competent *E. coli* directly without any further treatment. Fluorescent colonies were purified by restreaking and plasmids were obtained from fluorescent colony clones. These plasmids were subjected to DNA sequence analysis to confirm the nature of the fused ORF obtained.

**Figure 1.**
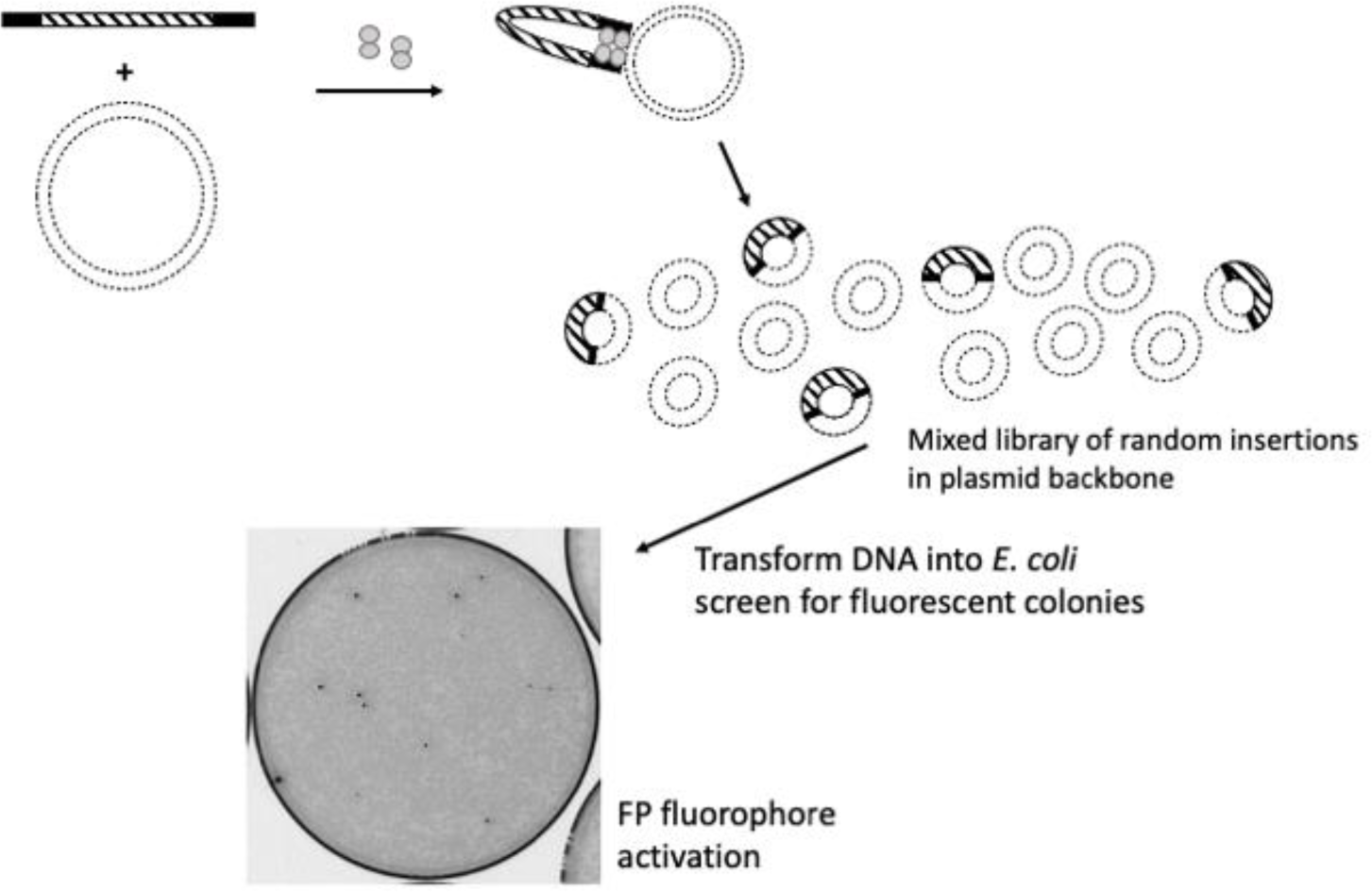
Tn5 Transposase (gray dumbbells) binds to FP ORF amplicons (black and hashed rectangles) at 19-nucleotide Tn5 binding sites (black). This complex interacts with target DNA (dashed circles) at random sites. Following transformation and incubation at 30°C fluorophore activation allows identification of fluorescent colonies (black spots, gray spots are non-fluorescent colonies).

Transformations were incubated according to the manufacturer’s instructions and 100 ul was plated onto each of 10 plates. The final volume of the transposition reaction was 11 ul, enough for 11 transformations resulting in 110 plates per transposition if desired. This procedure is designed to maximize the number of colonies screened. Transformations must be plated at high colony density (5×10^2^ per plate). Plates containing 1×10^2^ colonies per plate or less do not provide enough candidates to recover fluorescent colonies. This is why we employ MAX Efficiency DH5alpha competent cells, but any strain with comparable transformation efficiency can be used (>1 x 10^9^ transformants/µg plasmid DNA). It is essential to incubate the plates at 30 °C or less (room temperature works well). We have never obtained fluorescent colonies from 33 °C or 37 °C incubations. Furthermore, positive candidates identified at 30 °C that are re-grown at 37 °C will not be fluorescent. Fluorescent colonies were restruck and incubated at 30 °C to purify positive clones. Plasmid DNA from these cells was purified and employed to determine the site of the insertion of the FP ORF by DNA sequencing.

DNA sequencing was performed using a ‘universal’ sequencing primer strategy. The primer was designed to hybridize about 60-nucleotides from the 3’ end of the FP ORF and direct sequencing towards the 3’ end of the ORF, across the junction site and into the flanking target gene. Such a primer would allow confirmation of the presence of the FP, the maintenance of the correct reading frame at the junction, and the identification of where the insert is located within the target. We did not assess the 5’ junction because it must be correct for expression of a fluorescent product. DNA sequencing was performed by Quintara Biosciences, Cambridge, MA.

## Results and Discussion

The FP PCR amplicon was designed to contain no start codon, no stop codon and a single open reading frame from end to end. Therefore, in order for a FP to be expressed the FP ORF must be inserted in the correct orientation between adjacent codons to preserve the reading frame. The subsequent colony screen eliminates all insertions that land in the plasmid backbone, are in the incorrect orientation or are out frame insertions. The expected frequency of positive fluorescent colonies can be estimated. The manufacturer of EZ-Tn5 suggests that about 1 plasmid in 200 will contain a single insertion. The ratio of the target gene size to the total plasmid size influences the frequency (in our experiments with the EF G ORF this ratio is 2109bp/6837bp). The orientation of the inserted FP ORF relative to the transcription and translation of the target ORF was random so the frequency of a fluorescent positive colony was reduced by an additional one half. The probability of insertion between codons (not within a codon) must be accounted for (one out of three). Other factors that cannot easily be controlled for but can affect the frequency of positive colonies include the expression level of the fused ORF, and target protein structure (for example proteins containing multiple independent domains would be expected to offer more sites for successful insertion). By multiplying the factors for which there were numbers the expected positive colony frequency for the case of the EF G expression plasmid employed here was 1 positive colony for every 3890 screened. The observed frequency of in frame fusions (Supplemental Table S2-1) was much higher suggesting that for this ORF uncontrolled factors (expression level of the fused ORF, and target protein structure) significantly affected the frequency of obtaining in frame fusions and fluorescent colonies. In the case of the Era ORF (43% of the EF G ORF) in frame insertions occurred at more than 10-fold lower compared to EF G (Supplemental Table S2-2).

To demonstrate the simplicity and robustness of this method we employed it in an undergraduate laboratory class that included all biology majors with a very wide range of laboratory training. Between 2014 and 2020, 511 students in both spring and fall semesters undertook the experimental protocol detailed in the class lab manual (see File S1). To monitor robustness across multiple class sections in different semesters the undergraduate laboratory class module was conducted using only one target ORF (EF-4) and one transposable amplicon containing the GFPmut3 ORF. All class sections across all years successfully obtained fused ORFs for each student for further study. Figure 2 provides a map of all insertion sites documented between 2014 to 2020. Some domains (1 and 2) were particularly good targets for insertion whereas others were very rarely targeted (only a single insertion was ever observed in domain 4). This distribution of insertion sites would not be possible to predict by an *ab initio* approach highlighting the usefulness of the method; the ability to generate a very large library of useful fusion proteins.

**Figure 2.**
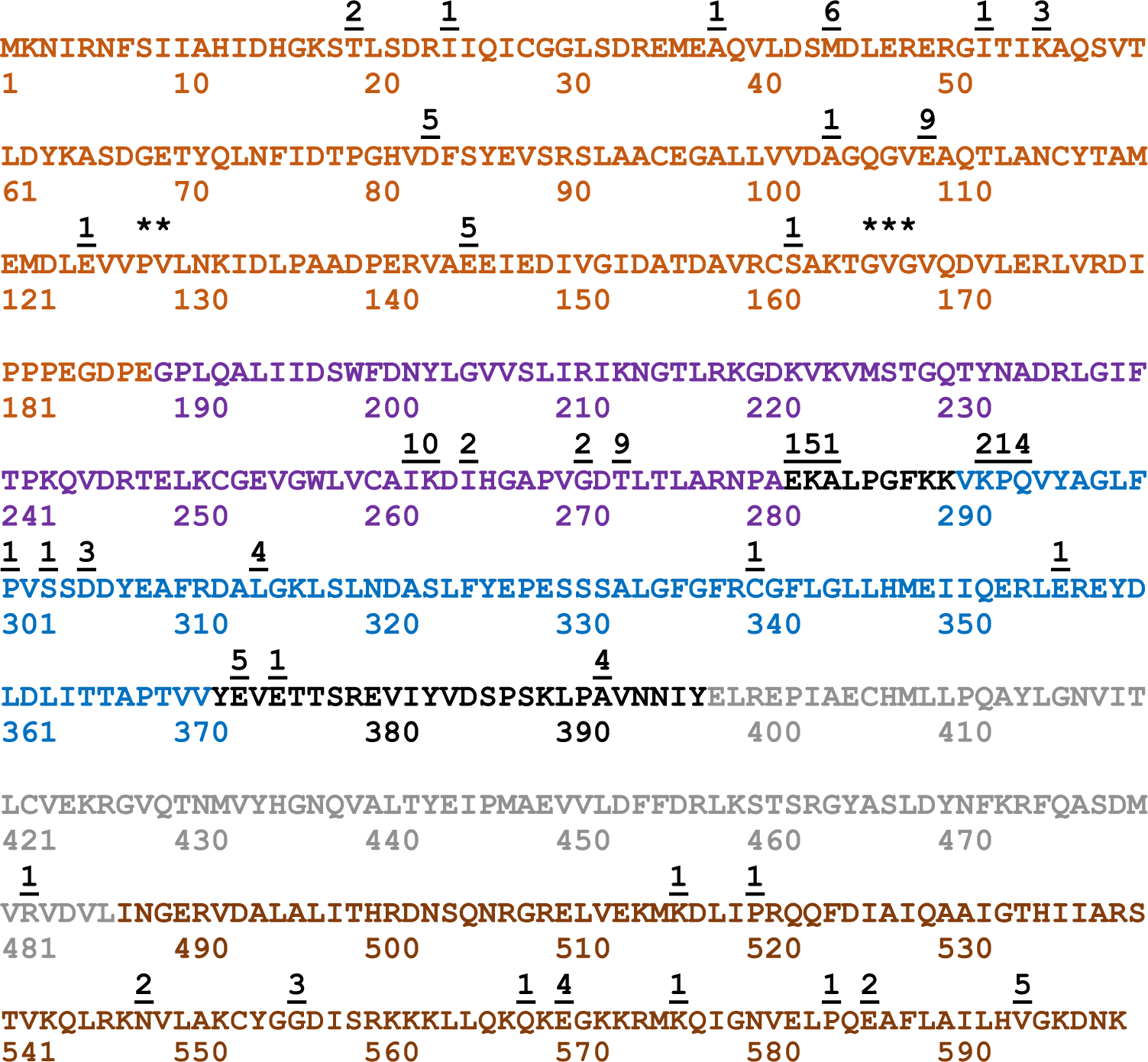
The cumulative data obtained from undergraduate laboratory class is shown. From 2014-2020 a single target ORF (EF 4) was employed in transposition reactions with the FP ORF GFPmut3. The amino acid sequence of the EF 4 ORF is shown. The colored amino acid residues indicate EF 4 domains: domain 1 (tan residues 1-188), domain 2 (purple residues 189-281), domain 3 (blue residues 291-371), domain 4 (gray residues 398-486), and the C-terminal disordered domain (brown residues 487-599). The underlined numbers above the sequence indicate insertion sites that were confirmed by student researchers and the number of times each insertion site was recovered. ** indicates the insertion at codon 128 was recovered 20 times and codon 129 twice. *** indicates a cluster of insertions at codons 167 (twice), 168 (13 times) and 169 (once). The cluster at 282, 283, and 284 was recovered once, five times and once respectively and at 292, 293 and 294, twice, once and four times respectively. These data were derived from 159 separate student experiments.

In parallel to demonstrate the broad applicability and potential of the method investigations were undertaken to explore the effect of different FP ORFs, and the effect of the linker sequence between the target and the FP ORF. Finally, the approach was also used on several different target ORFs. Three specific questions have been addressed: (a) does alteration of the sequence that flanks the FP ORF affect the efficacy of successful transposition, (b) can other FPs replace the GFPmut3 allele, (c) is it possible to apply MORFIN to other target genes?

The sequence that flanks the FP ORF must include the DNA sequence required for Tn5 transposition, but additional codons can be included in linker sequences. EZ-Tn5 requires a specific 19-nucleotide inverted repeat at the 5’ and 3’ ends of a DNA sequence destined for transposition (7). The 19-nucleotide sequence represents a binding site for the Tn5 transposase (Figure 1). Insertion into target DNA results from a double strand cleavage which is staggered with a nine-nucleotide overhang. Due to this, three codons at the 3’ end of a cleavage site are a repeat of the same codons found at the 5’ end (Figure 3). Therefore the 5’ flanking sequence would contain six spacer codons between the target protein ORF and the FP ORF, whereas as the 3’ flanking sequence would contain 9 extra codons (Figure 3). Although the 19-nucleotide recognition sequence cannot be altered it is possible to insert additional codons between this sequence and the FP ORF. Investigations into the optimal lengths of linker sequences between protein domains and the composition of those sequences guided our initial experiments (8,9). Based on these data we created a linker to insert 3 extra codons at the 5’ end to balance the number of codons at each end to nine extra codons. In addition, linkers were created to include 12 extra codons at each end (Figure 4, Table 1).

**Figure 3.**
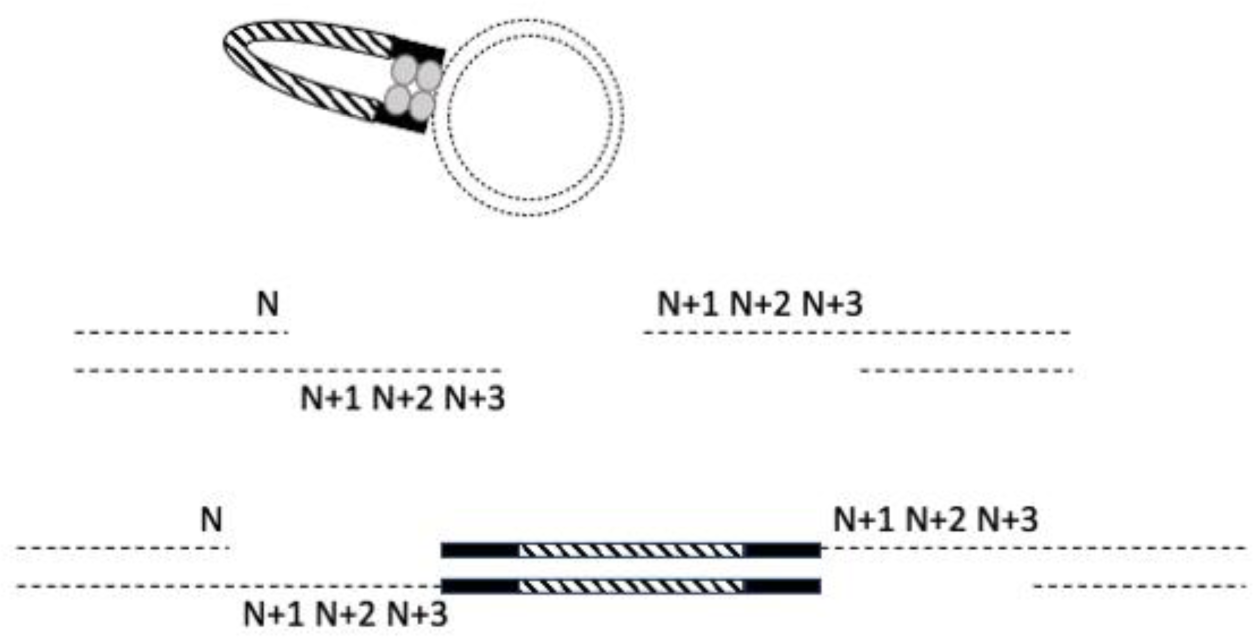
Tn5 Transposase (gray dumbbell color) binds randomly to the target DNA (dashed circle) and cuts target DNA strands at sites nine nucleotides apart generating a nine-nucleotide complimentary overhang. The transposase ligates the inserted DNA generating a double strand structure with 5’ and 3’ nine-nucleotide gaps. The gaps in the resultant double strand circle are filled in by host DNA repair polymerases following transformation. Codon N is the codon that immediately precedes the Tn5 cut site. Codon N+1, N+2, and N+3 represents the three codons following N which comprise the nine-nucleotide overhang. The result of ligation by Tn5 and host cell repair is that codons N+1, N+2, and N+3 were repeated on the 3’ side of the FP ORF (hashed rectangle). The 19-nucleotide recognition sequence for Tn5 is represented by the black rectangle. Only cleavages between codons are recovered because cleavage within a codon produces an out of frame ORF. Out of frame ORF fusions fail the colony screen for fluorescence.

**Figure 4.**
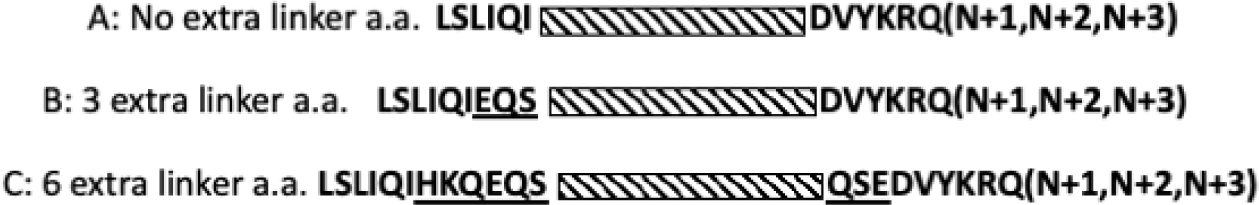
Testing effects of altering linker sequence (black bold capital letters) between the target ORF and the FP ORF (hashed rectangle). (A) The 19-nucleotide Tn5 recognition element is a defined sequence that must always be present at the 5’ and 3’ ends of the inserted DNA as inverted repeats. The first 18 nucleotides would encode the amino acid sequence **LSLIQI** and the 19^th^ nucleotide would become the first base of the first codon of the FP ORF. At the 3’ end the first nucleotide of the inverted repeat would replace the third nucleotide of the ORF’s stop codon creating a read through ORF. The subsequent 18 nucleotides encode the amino acids **DVYKRQ**. Due to the staggered cut of the Tn5 transposase codons N+1, N+2, N+3 were repeated at the 3’ end of an insertion (Figure 3) comprising additional non-native sequence. (B) To make the number of 5’ and 3’ inserted codons symmetrical three additional codons were inserted at the 5’ end (underlined letters represent the amino acids encoded by this modification). (C) To increase the length of the linker region to optimal 12 codons and insert codons encoding more favorable linker amino acids, codons were added that would encode the underlined amino acids. George and Heringa (8) and Suyama and Ohara (9) were used as a guide to design optimal linker length and favorable inter-domain amino acid linker sequence. PCR primers employed to insert these modifications are listed in Table 1.

This linker set was employed to create amplicons derived from the open reading frames of three different FPs, GFPmut3, mCherry2, and mYPET (Figure 5). This panel of amplicons was used in initial experiments to examine effects associated with altering either the linker sequence or the FP ORF.

**Figure 5.**
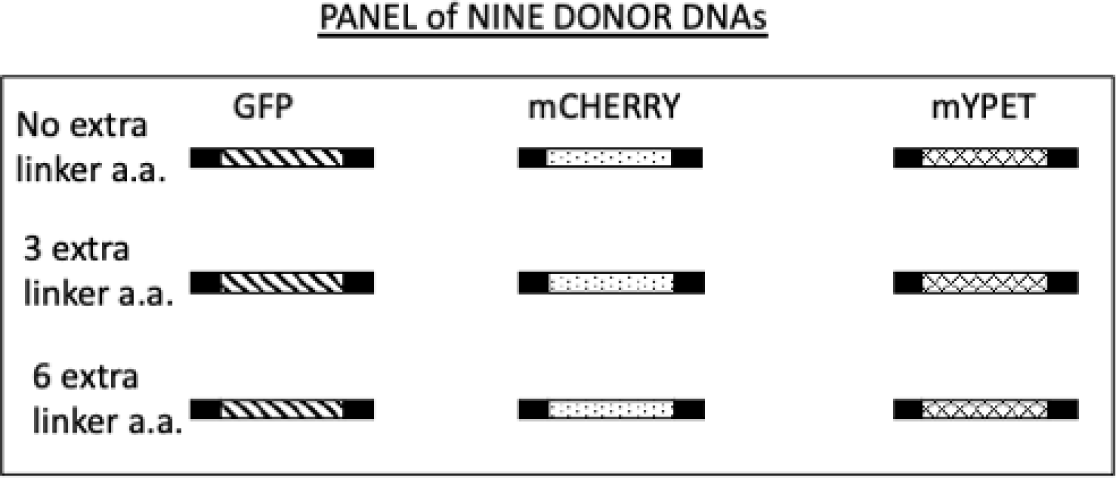
Schematic of donor amplicons generated as substrates for MORFIN mutagenesis. The black rectangle indicates the position of 5’ and 3’ linker sequence. ’No extra linker a.a.’, ‘3 extra linker a.a.’ and ‘6 extra linker a.a.’ are explained in Figure 4. The diagonal filled rectangle represents the ORF for GFPmut3, the dotted rectangle represents the ORF for mCherry2, and the hashed rectangle represents the ORF for mYPET.

Table 2 lists the ORFs that were tested using the amplicons created. It was beyond the scope of this study to test the entire panel of amplicons in all the ORFs listed in Table 2. The reason for this is that nine transpositions are required to test the entire collection of amplicons. Each plate screen requires a minimum of ten plates to obtain a collection of fluorescent positive colonies (often more than 10). Accounting for different frequencies of positive colonies, testing the entire panel would require about 10^3^ plates. The EF G ORF, the FtsZ ORF and the MreB ORF were tested using the entire panel of amplicons. The EF G ORF yielded fluorescent colonies and in frame fusions from all combinations of linkers and FPs except one combination: mYPET and no extra linker amino acids (Figure 6). In fact, we never obtained any positive candidates from that combination, suggesting that the mYPET FP is particularly sensitive to flanking sequence. Additional evidence of the linker spacer effect on mYPET fusions is apparent in Figure 6. When mYPET was combined with 6 extra a.a. linkers fluorescent colonies were observed of varying signal intensity. A more intense signal would be either a consequence of lower fusion protein turnover, or a higher percentage of correctly folded expressed protein, or both. The mYPET 6 extra a.a. linker also allowed for fusion proteins to be obtained for ORFs that were otherwise recalcitrant to the MORFIN approach. The FtsZ and MreB ORFs only gave rise to positive candidates with in frame fusions with one combination: mYPET with six extra linker amino acids (Figure 7). Using homology modeling of MreB Bendezu et al (10) identified a surface loop centered around codon 228 and employed site directed mutagenesis to create a functional internal fusion to mCherry. Among the MreB candidates from this work we identified mYPET fusions after codon 228 and nearby at 235. Site directed mutagenesis was employed to investigate internal FP fusions to FtsZ (11). From their collection one fully functional insertion was obtained between FtsZ codon 55 and 56. In this study an insertion was obtained after codon 57.

**Figure 6.**
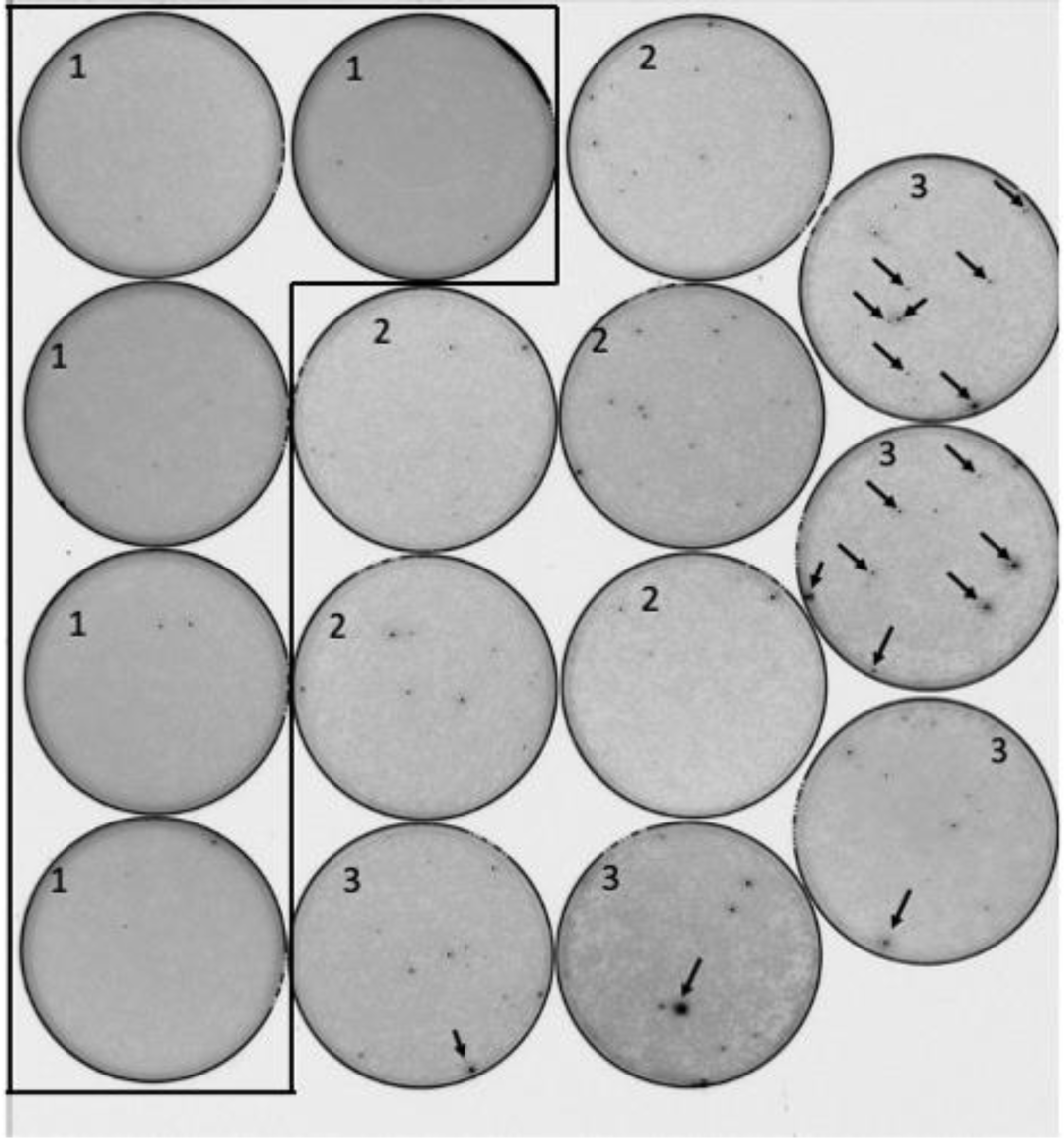
The unexpected sensitivity of mYPET to linker sequence. Transposition reactions were transformed directly into *E. coli* and plated at high colony density. After a 36-hour incubation at 30°C the plates were imaged using a laser plate scanner. Non-fluorescent colonies are light gray, fluorescent colonies are black. Examples of prominent black fluorescent colonies are indicated by the arrowheads. The results from three different amplicons are shown: (1) mYPET with no extra linker, (2) mYPET with 3 extra linkers, (3) mYPET with 6 extra linkers. The meaning of ‘no extra linker’, ‘3 extra linkers’, and ‘6 extra linkers’ is explained in Figure 3. The target ORF in this case expressed EF G. All black colonies were restruck to purify clones that produced fluorescent proteins. Purified clones were subjected to plasmid purification and pure plasmids were sequenced to confirm insertion sites. The tiny black specks found on plates numbered 1 were not fluorescent bacterial colonies.

**Figure 7.**
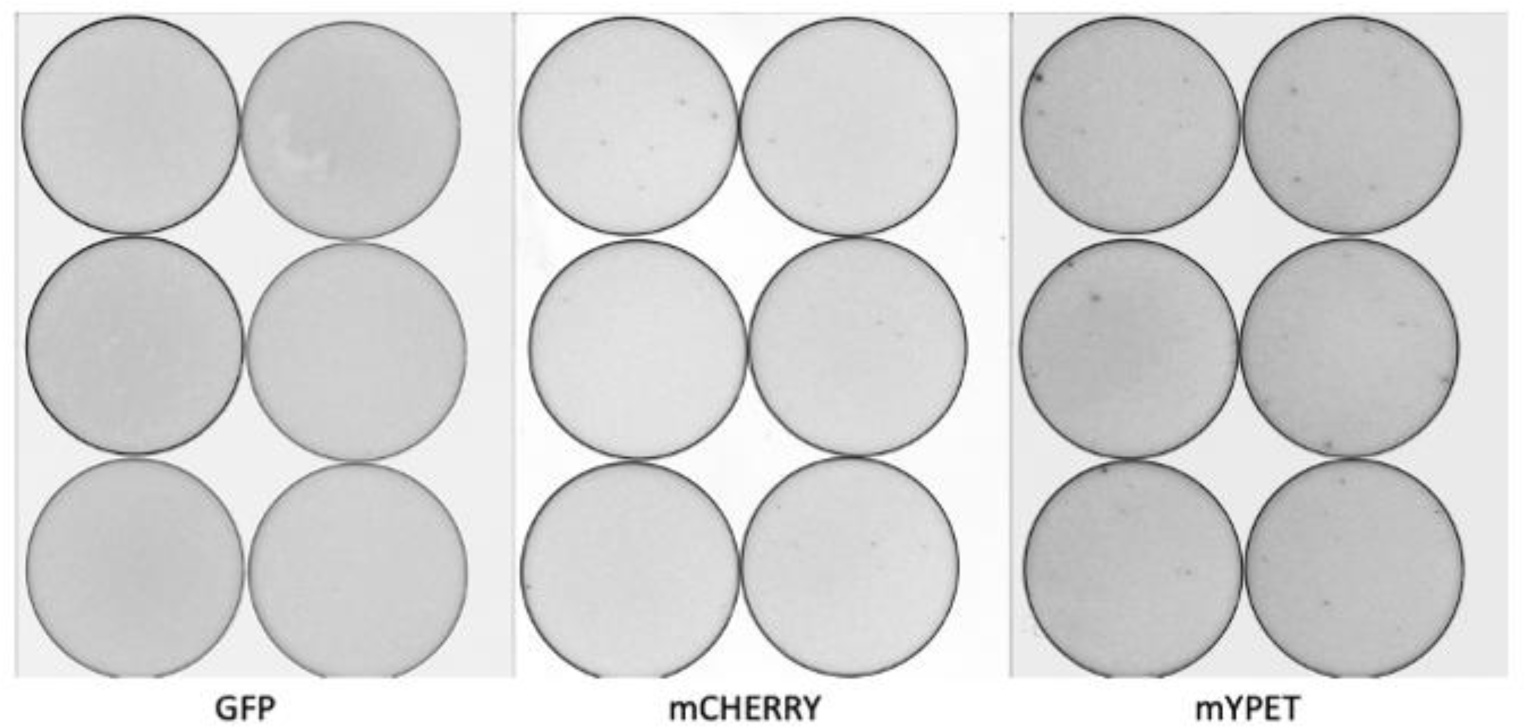
An example of the restrictive nature of the FtsZ ORF. Transposition reactions were transformed directly into *E. coli* and plated at high colony density. After a 36-hour incubation at 30°C the plates were imaged using a laser plate scanner. Non-fluorescent colonies are light gray, fluorescent colonies are black. The results from three different fluorescent proteins are shown. In each case the linker was the 6 extra linker (Figure 3). Fluorescent colonies were only found when mCherry or mYPET was employed as the FP ORF. All of the mCherry candidates were off target. In frame FP – FtsZ fusions were only obtained from mYPET transposition reactions. The same result was obtained for the MreB ORF. All black colonies were restruck to purify clones that produced fluorescent proteins. Purified clones were subjected to plasmid purification and pure plasmids were sequenced to confirm insertion sites.

It is important to consider protein production levels because if an in frame fusion is expressed at a low level or is rapidly turned over it would fail to be detected in the colony screen. FtsZ presents a particular challenge because expression levels are tightly regulated in dividing cells. For this reason, a plasmid backbone with a lower copy number was used (Table 2). The antibiotic marker gene on plasmids expressing the FtsZ and MreB ORFs was CAM^R^ conferring resistance to chloramphenicol. Although the CAT ORF was not designed as a target in these studies in frame fusions to the chloramphenicol acetyltransferase ORF (CAT) were obtained in the fluorescent colony screen (one after codons 20, 208, and 214, and nine after codon 6). These were off target, but informative in frame fusions. They were confirmed by plasmid purification, retransformation, and DNA sequencing. Structural analysis of insertions sites showed that the insertions were located at the external face of monomers and could be accommodated within a homo-trimer structure. These fusions were both fluorescent and conferred resistance to chloramphenicol confirming that both partners of the fused protein were functional. The CAT ORF is only 219 codons, so it represents a small target. In addition, the functional CAT protein must assemble into a homotrimer (12). It was not expected that a FP ORF (with the number of codons similar to the target ORF) could be inserted into the CAT ORF and that a functional trimer could assemble. This observation strongly supports the broad application for this technology and its potential to recover fully functional protein chimeras; however, the gene encoding CAT should not be employed as a selectable plasmid marker for the MORFIN approach because off – target insertions were not rare. In frame fusions were not ever seen within the Beta-lactamase ORF (this observation includes hundreds of fusions examined in the undergraduate laboratory classes). Beta-lactamase is a secreted protein, and it is not a surprise that proteins that cross a membrane would not easily accommodate a FP fusion with a functioning fluorophore. ORF expression plasmids using Amp^R^ as the selectable marker are preferable in the MORFIN approach.

A consideration regarding the target ORF is that the frequency of obtaining positive FP-target protein fusions depends on the size of the target ORF and the domain structure of the target protein. In an effort to investigate the affect of domain structure on insertion frequency a panel of related GTP-binding proteins were tested (Table 2; EF G, EF 4, EngA, Era). Each of these four ORFs were present on the same plasmid backbone and targeted by the FP amplicon containing the FP mGFPmut3 with no added linker a.a. EF G, like EF 4, is a large multi-domain protein and gave rise to a positive ORF insertion frequency of 70 inserts/40,000 transformants screened (Supplement Table S2-1. The EF G ORF is 703 codons, and the protein contains six distinctly folded domains (13). The EF 4 ORF is 599 codons, and the protein contains 5 domains (14). EngA and Era (Table 2) are related GTPases whose ORFs contain three and two domains respectively (15, 16). The Era ORF contains 301 codons and was found in fluorescent fusions ten time less frequently than EF G. The EngA ORF contains 490 codons, and it was found at frequency about five times higher than Era (Table S2-1, S2-2 and S2-3). One recurring observation across all four ORFs was that the GTP-binding domain was frequently a target for in frame FP insertions so this method would be useful to explore biochemical and cellular functions of GTP-binding proteins.

One important consideration in selecting a fluorophore protein is that the mYPET ORF gives rise to a fluorescent protein that must be detected by laser activation. This means that access to expensive detection devices (such as a laser plate scanner) is necessary. The mGFPmut3 ORF product can be detected with inexpensive handheld UV devices. Colonies that express mCherry2 can be identified because they turn red even in ambient room light. These considerations may be critical for implementation into undergraduate curricula and research.

In conclusion the MORFIN method is a simple and powerful research tool. Transformation of transposition reactions with no intervening steps eliminates time and resource intensive steps traditionally employed in methods to create fluorescent protein fusions (11, 17, 18, 19). The power of the method arises from subsequent fluorescent colony screen. The transposed DNA must land within an ORF, in the correct orientation, reading frame, and be expressed highly enough for a fluorescent signal to arise. The fluorophore of fluorescent proteins is exquisitely sensitive to the structure of the protein (20). Since the colonies are fluorescent the fluorophore must be precisely and correctly folded. It is extremely likely that the surrounding target protein is not misfolded because prior research demonstrates that GFP fluorescence is negatively impacted when it is flanked by misfolded sequence (21–27). Correct folding is supported by the evidence presented here that fully functional fusions in the CAT ORF were obtained. Furthermore, the plate screen apparently strongly screens out fused ORFs that would disrupt protein structure. In all cases where detailed structural mapping has been done (EF G, EF 4, Era and CAT ORFs) all of the insertions are localized at external surfaces of the protein structure. The activities described here are perfectly suited to provide students with experience in a variety of commonly used molecular biology methods. The module presented in the laboratory manual (see File S1) is designed to give students ownership in the project by providing each student with a random, unique insertion site to characterize. Students must utilize critical thinking skills to develop appropriate predictions of the results and to interpret the data from the experiments. MORFIN can additionally be used to quickly create a large, mixed library of random mutations to screen for promising candidates that can be investigated in a continuing research program. Although we only tested ORFs derived from *E. coli* genes we expect MORFIN to be applicable to ORFs from any source if the ORF’s product can be expressed in the *E. coli* cytoplasm.

## Data Availability

The raw data from all of our FP fusions are available.

## Acknowledgements

We would like to thank all the Emmanuel College students, faculty, and staff who have participated in this learning module. We would like to thank Tom Bernhardt and Nick T. Peters, Harvard Medical School (N.T.P. current affiliation, Microbiology Undergraduate Program, Iowa State University) for discussions, research support and access to a laser plate scanner.

## File S1. Supplementary Material

**Class Lab Manual Step – by – Step Protocols Using the MORFIN Method**

**Table of Contents**

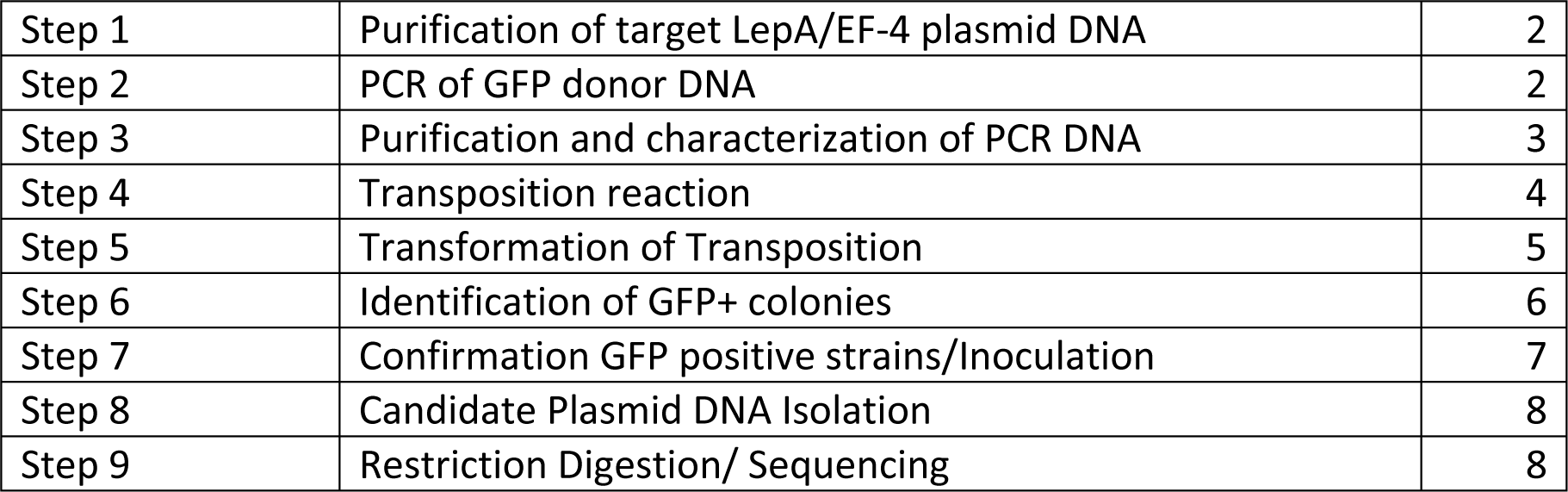

### Experimentation Schedule

#### Overview

Initially the target plasmid (pPEM109) containing the *lepA* gene will be isolated and donor transposon DNA will be synthesized. After characterizing these pieces of DNA, they will be combined with Tn5 transposase. This reaction will generate an insertion library as illustrated in FIGURE 1. This library will be transformed into *E. coli* in order to isolate insertion-positive candidates by screening for fluorescent colonies growing on plates. The plasmid derived from fluorescent colonies will be purified and compared to the original target DNA. The characterization will include determining the relative sizes of the plasmid DNA, and the EF-4 protein produced, DNA sequence analysis, and western blotting to prove that the insertion-positive candidate contains GFP sequence. DNA sequence information will be used, in conjunction with information about the three-dimensional protein structure to determine where the GFP sequence is inserted within the structure of EF-4. Functional assays will be performed to assess whether the newly created GFP fusion proteins are active.

## STEP 1: PURIFICATION OF TARGET LEPA/EF-4 PLASMID DNA

### Purpose

To amplilfy and purify plasmid pPEM109, containing the *lepA* gene as a GFP target, for an *in vitro* Tn 5 transposase-catalyzed transposition reaction

### Procedure

Plasmid Purification.

### Each student will start with 1.5 mL of cultured *E. coli>* containing plasmid pPEM109

1. Obtain a microfuge tube containing 1.5 mL of saturated overnight *E. coli* culture (containing desired plasmid pPEM109). Label top of tube with your initials.
2. Purify the plasmid using an appropriate miniprep or similar kit according to the manufacturer’s instructions.
3. The DNA will be stored at -20^°^C until the next lab meeting.

## STEP 2: GENERATION OF TRANSPOSITION REACTION “DONOR” DNA USING PCR

### Purpose

To use the Polymerase Chain Reaction (PCR) to create a linear double-stranded piece of DNA that can serve as the “donor” DNA in an *in vitro* Tn5 transposase-catalyzed transposition reaction.

### Procedure

1. Each student should label a single PCR tube with their individual initials. *Note: The PCR tubes are different than microfuge tubes. They are smaller to better fit into the thermocycler machine and have have thinner walls more conducive for heat conduction*.
2. Add the reagents to <u>PCR tubes on ice in the order shown below</u> to prepare the following 50 uL reactions: HINT: Put a checkmark next to protocol reagents “Reagent added?” as you add them to help you remember what has been added to each tube.

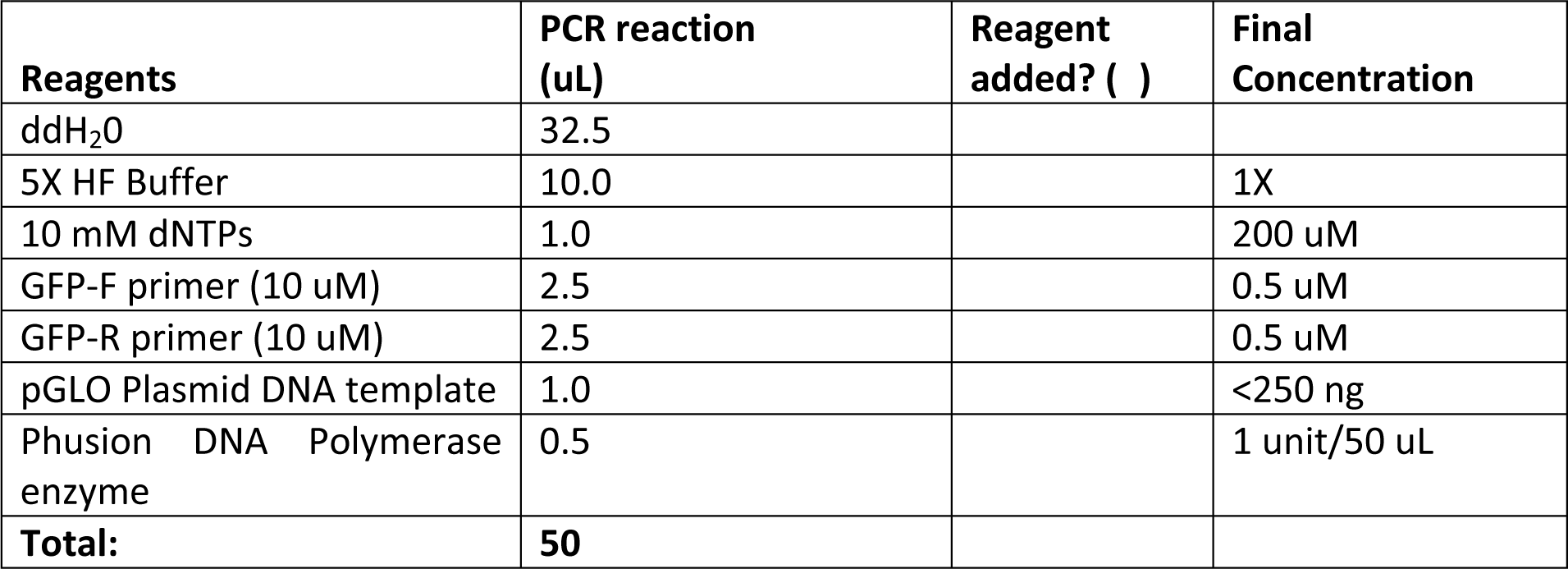
3. Seal the PCR reaction tubes with caps, making sure they are tightly closed. Mix the reaction by tapping the side of the tube. Spin down in a mini microcentrifuge to collect contents.
4. Keep your reactions on ice until ready to load into the PCR machine that has been programmed as shown below:

#### Cycling conditions for GFP PCR

1. 98°C, 30 sec.
2. 98°C, 10 sec.
3. 62°C, 20 sec
4. 72°C, 30 sec
5. Repeat 2-4, 30X
6. 72°C, 5 min.
7. 10°C, Hold

## STEP 3: PURIFICATION & CHARACTERIZATION OF PCR PRODUCT (“DONOR DNA”)

## To check that the PCR product was produced, purify the PCR product, and determine the concentration of the PCR product

### Procedure

#### PCR Clean-up

1. Clean up the plasmid using PCR cleanup kit according to the manufacturer’s instructions.

#### Nanodrop

Use 2 ul of your cleaned up PCR sample to determine the DNA concentration using a Nano-Drop spectrophotometer. Your lab instructor will demonstrate how to use the Nano-Drop spectrophotometer. Record this value in your lab notebook, including concentration (ng/uL) and 260/280 purity ratio.

#### Agarose Gel Electrophoresis

1. Remove 5 uL of your cleaned-up PCR reaction and add into a new microcentrifuge tube along with 3 uL of loading dye. Pipette up and down to mix. **<u>Note: Do not add dye to your entire PCR</u> <u>SAMPLE</u>**!
2. Load the 8 uL sample into a 1% agarose gel containing fluorescent dye and perform electrophoresis. Save the remainder of the cleaned-up PCR for future use.

## DATA ANALYSIS

1. Were you successful in amplifying the desired PCR product? How can you tell? Be sure to describe any expected bands and relative sizes.
2. Compare the yield of your reaction compared to your partner’s reaction. What could account for any differences that you observe?
3. What sequences are contained within your PCR product, and how will the PCR DNA be used in future experiments?
4. Consider any potential sources of error in the experiment (even if not observed) and how that may have affected the outcome.

## STEP 4: TRANSPOSITION REACTION

## To set up the transposition reaction with GFP PCR donor DNA and pPEM109 plasmid DNA

### Procedure

#### Transposition Reaction

Tn5 transposase is expensive, but one 10 microliter reaction is enough for 8 pairs of lab partners. The reaction will be set up by the instructor and shared with students.

1. The transposition reaction requires 200 ng of plasmid pPEM109 (target DNA) and a 1:1 molar ratio of donor transposon (source of the GFP gene). The students will have to calculate the volume of plasmid pPEM109 DNA, GFP DNA, and water that are required. Then, set up the transcription reaction as described in the table below

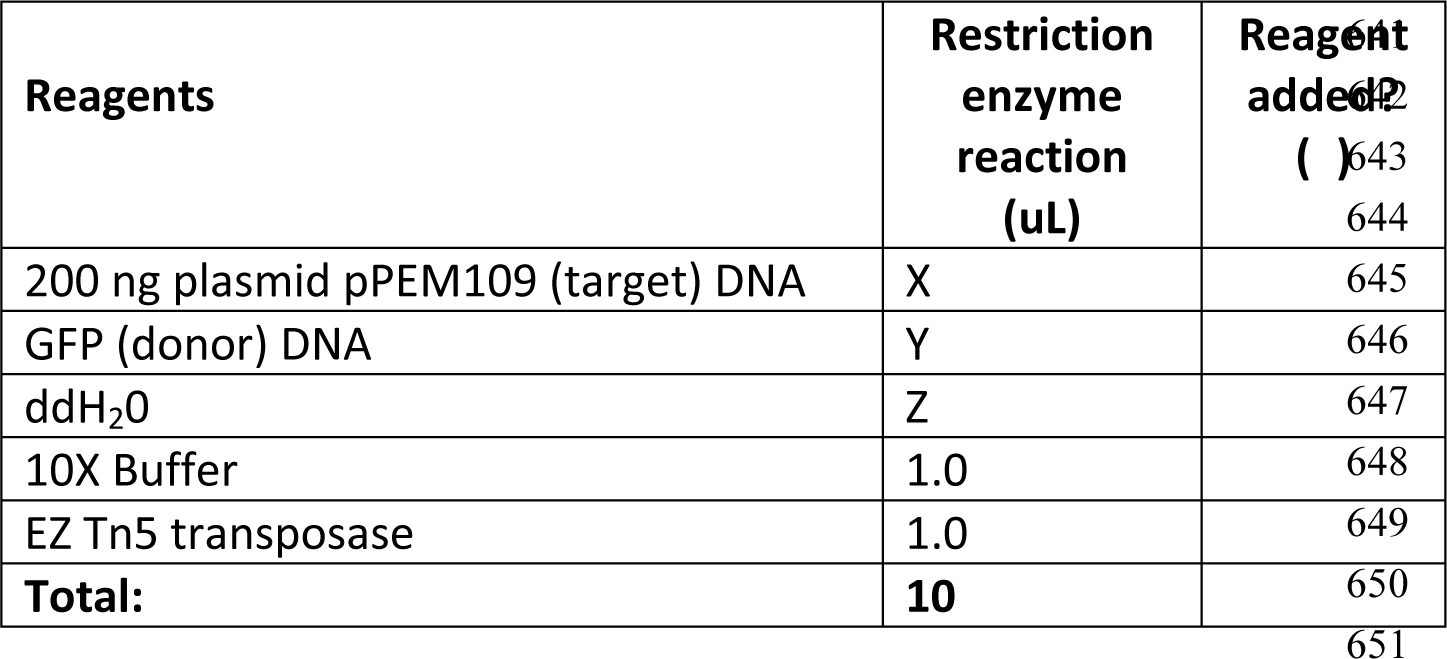
2. Incubate the reaction for 2 hours at 37°C.
3. Terminate the reaction by adding one microliter of 10X stop solution and incubating for 10 minutes at 70°C.

At this stage the sample can be stored at -20 degrees C. Each lab partner pair will use one microliter of this reaction mixture in STEP 5 to transform ultra-competent *E. coli* cells.

## STEP 5: TRANSFORMATION OF TRANSPOSITION MIXTURE INTO *E. coli*

### Purpose

To introduce a mixed library of GFP insertion plasmids into bacteria to measure GFP fluorescence *in vivo*

### Procedure

#### Important note

Correct handling of High Efficiency Competent cells is crucial to obtain the maximum number of transformants. Cells must be removed from -80 freezer and placed directly in ice. Cells must be thawed gently on ice. Maintain ice temperature until heat shock. For heat shock be exact with the incubation time and be sure cells are transferred from ice directly to 42 degrees. Do not alter incubation times.

1. Thaw *E. coli* strain DH5alpha-F’lacIQ in ice bucket.
2. Label a sterile 1.5 mL tube with your initials and “+library”. Add 1 ul of transposition reaction mixture into the labeled tube and place on ice.
3. Observe the *E. coli* DH5alpha-F’lacIQ cells and give the tube a quick flick to be sure cell suspension is uniform. Tap tube on bench to collect cells and place back on ice. DO NOT ALLOW CELLS TO WARM UP DURING TRANSFER. Try to be precise and quick with your transfer. Add 100 ul of thawed cells to the tube containing 1 ul of transposition library. **NOTE:** AT THIS POINT IN THE PROCEDURE IT IS CRITICAL THAT SAMPLES BE MAINTAINED ON ICE.
4. Incubate your sample tube (containing 1 ul of transposition mixture and 100 ul of cells) on ice for 30 min.
5. Heat shock cells by transferring sample tube from ice directly into a 42°C heating block. Incubate for EXACTLY 45 seconds.
6. Transfer sample tube directly from heating block back to ice bucket. Incubate 2 minutes on ice.
7. Add 0.9 mL of SOC medium (maintained at room temperature) to the cell mixture.
8. Place sample tube in a rotator in a 37°C incubator for 1 hour.
9. Spread cells from sample tube onto LB+Amp plates by aliquoting 0.1 mL onto each plate and spreading with sterile beads. You should generate 10 plates from this step. Plates should be labeled with group initials and the date.
10. Incubate plates <u>in a lab drawer (at room temperature)</u> until next meeting.

***Note:*** *4 days at room temperature will yield colonies but they may be small depending on ambient temperature (20-25°C). Switch plates to 30°C for final overnight incubation the day prior to next lab meeting*.

## STEP 6: IDENTIFICATION AND RE-STREAKING OF GFP POSITIVE COLONIES

## To identify GFP positive transformant colonies and obtain genetically pure isolates for additional characterization

### Procedure

1. Inspect plates using hand held UV light, carefully inspecting all colonies for signs of fluorescence. Circle the back of the plate with a marker if GFP positive colonies are identified.
2. Record number of <u>ALL</u> green glowing colonies. Record total number of colonies by counting section and estimating for remainder of plate. Estimate frequency of GFP+ colonies over total transformants. If you have no colonies record that. If you have a confluent lawn of cells record that as a “lawn”. Your instructor will assist you with this.
3. <u>Each student should re-streak one GFP positive colony</u> onto a new LB Amp plate. If you obtained more than one green colony, pick the strongest GFP positive colony that is well separated from neighboring colonies. If you have no positive colonies borrow colonies from colleagues that have extra. Carefully pick GFP positive colonies, as instructed. Consult instructor about discarding transformation plates.
4. LABEL EACH ISOLATE WITH A DISTINCT NUMBER AND RECORD YOUR LABEL.
5. Incubate re-streak plates at room temperature until next meeting.

*Note to instructor: Keep several plates that have multiple GFP-positive colonies as a backup in case the re-streaked plates don’t grow properly*.

## DATA ANALYSIS

1. Was the transformation of the mixed plasmid library successful? Why or why not?
2. What are 2 different transposition events that could result in a GFP positive colony? What are 2 different transposition events that could result in a GFP negative colony?
3. Based on data up to this point, can you tell if the transposition and transformation was successful in generating a successful *lepA* gene-GFP gene ORF fusion strain? Why or why not?
4. Consider any potential sources of error in the experiment (even if not observed) and how that may have affected the outcome.

## STEP 7: CONFIRMATION OF GFP POSITIVE STRAINS AND INOCULATION OF POSITIVE CANDIDATES INTO LIQUID MEDIA

### Purpose

To confirm we have genetically pure GFP positive isolates and start liquid cultures for future analysis of plasmid DNA sequence and GFP protein expression

### Procedure

1. Lab partner pairs should have two plates from STEP 6. Each plate contains a re-streak of an original colony that was GFP-positive. Inspect plates to confirm GFP-positive colonies.
2. Record whether there is growth on each plate and record whether or not there are GFP-positive colonies. Consult with an instructor to determine your official candidate ID number (each student will have their own).
3. <u>Each student</u> pick one GFP positive colony using a sterile loop. If there is growth on both plates from, be sure that one colony is taken from each plate. If there is growth on only one plate or no growth on either plate, then consult with instructor to formulate a plan). Each GFP-positive colony should be transferred into one 15 mL sterile conical tube containing 1.0 mL of LB Amp. Incubate this tube at 30^°^C during the lab.
4. Incubate the conical tubes at 30°C overnight and arrange with your instructor to transfer these cultures to 4°C the following morning.

SAVE ALL GOOD RESTREAK PLATES IN CASE WE HAVE TO GO BACK TO THEM. LABELING IS IMPORTANT!

## STEP 8: CANDIDATE PLASMID DNA ISOLATION

### Purpose

To isolate pPEM109+GFP plasmid and total protein from liquid cultures containing the GFP-positive transformant

### Procedure

#### PLASMID PURIFICATION

1. Vortex to resuspend the cells from your 3 mL culture from STEP 7.
2. Transfer 1.5 mL of culture into a new 1.5 mL centrifuge tube. Label this tube “plasmid DNA” with your candidate number.
3. Use this sample to isolate plasmid DNA according to the STEP 1 procedure described in this manual. Store samples as -20°C.

## STEP 9: RESTRICTION DIGESTION OF PLASMID DNA. AGAROSE ELECTROPHORESIS; DNA SEQUENCING REACTION

### Purpose

#### To determine size of the GFP-positive plasmid <u>Restriction Digestion Set-Up</u>

1. Label a 1.5 mL microfuge tube with your candidate number and “pPEM109-GFP +Xba1” and proceed to set up restriction digestion as described in the table below: HINT: Put a checkmark next to protocol reagents “Reagent added?**”** as you add them to help you remember what has been added to each tube.

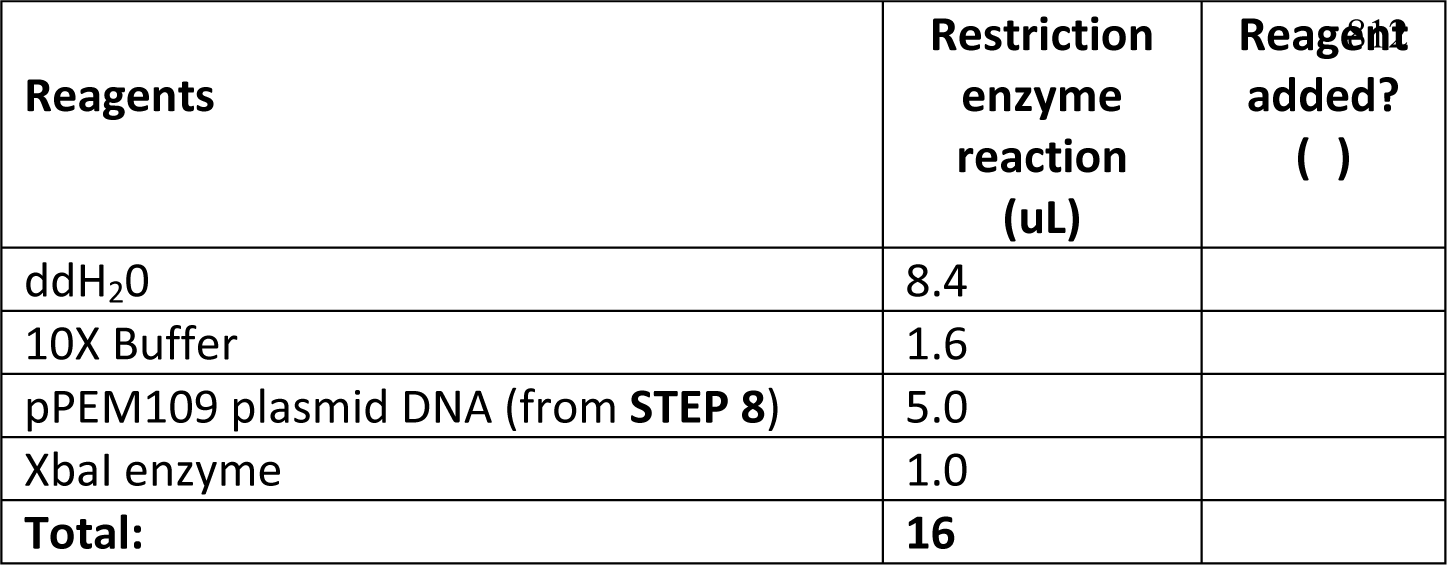
2. Label a 1.5 mL microfuge tube with your candidate number and “pPEM109 +Xba1” and set up:

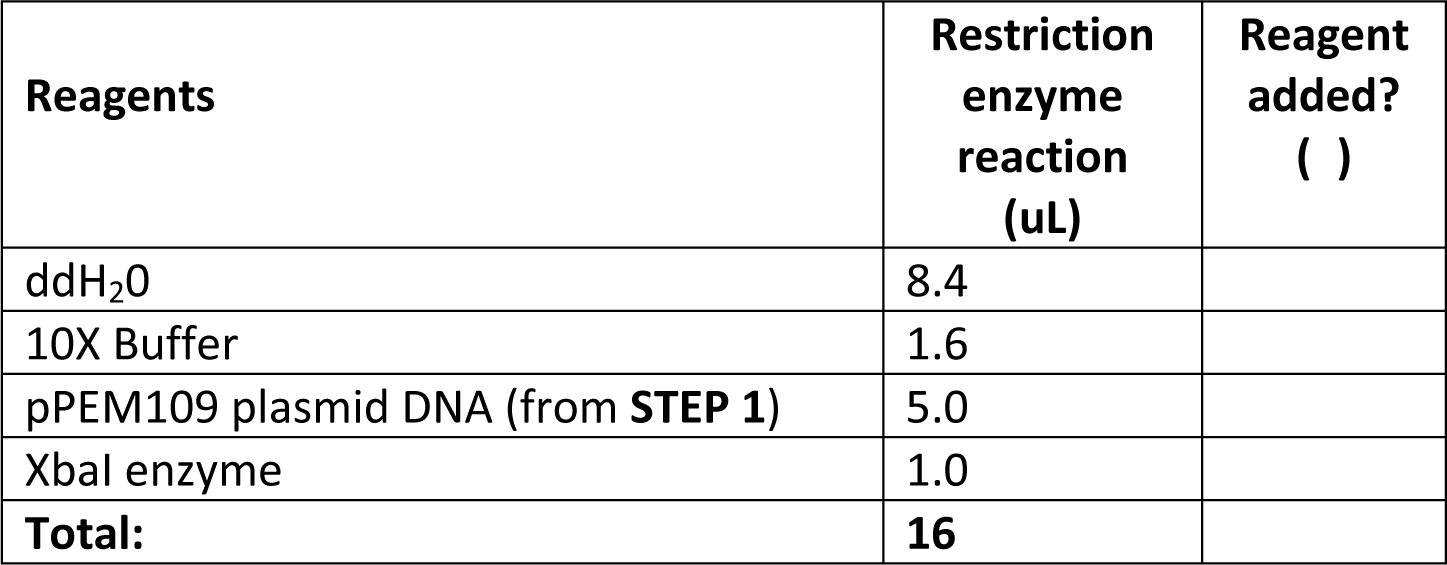
3. Incubate digestions at 37°C for 1 h. Add 4 uL of DNA sample loading dye to stop reaction.

Why is it important to compare STEP 1 results with STEP 8? What do you expect and why?

#### DNA Sequence Analysis

Remove 12 ul of STEP 8 plasmid DNA and place it in a PCR striptube tube containing 3 ul of GFP DNA sequencing primer (5 uM) as directed by your instructor. DNA sequencing samples will be sent to an outside sequencing lab for sequence determination. We will analyze these results when the sequence becomes available.

#### DNA gel electrophoresis

1. Load 8 ul of XbaI-digested DNA from both the STEP 8 and the STEP 1 DNA samples. Each student should have 2 samples to load onto the gel. Depending on gel lane availability, your instructor may also have you load an undigested control DNA sample. Run the gel as described in STEP 3. When the gel finishes, image the gels using BIO-RAD GelDoc.

**File S2. Supplementary Material: Tables S2-1, S2-2, and S2-3**

**Table S2-1.**
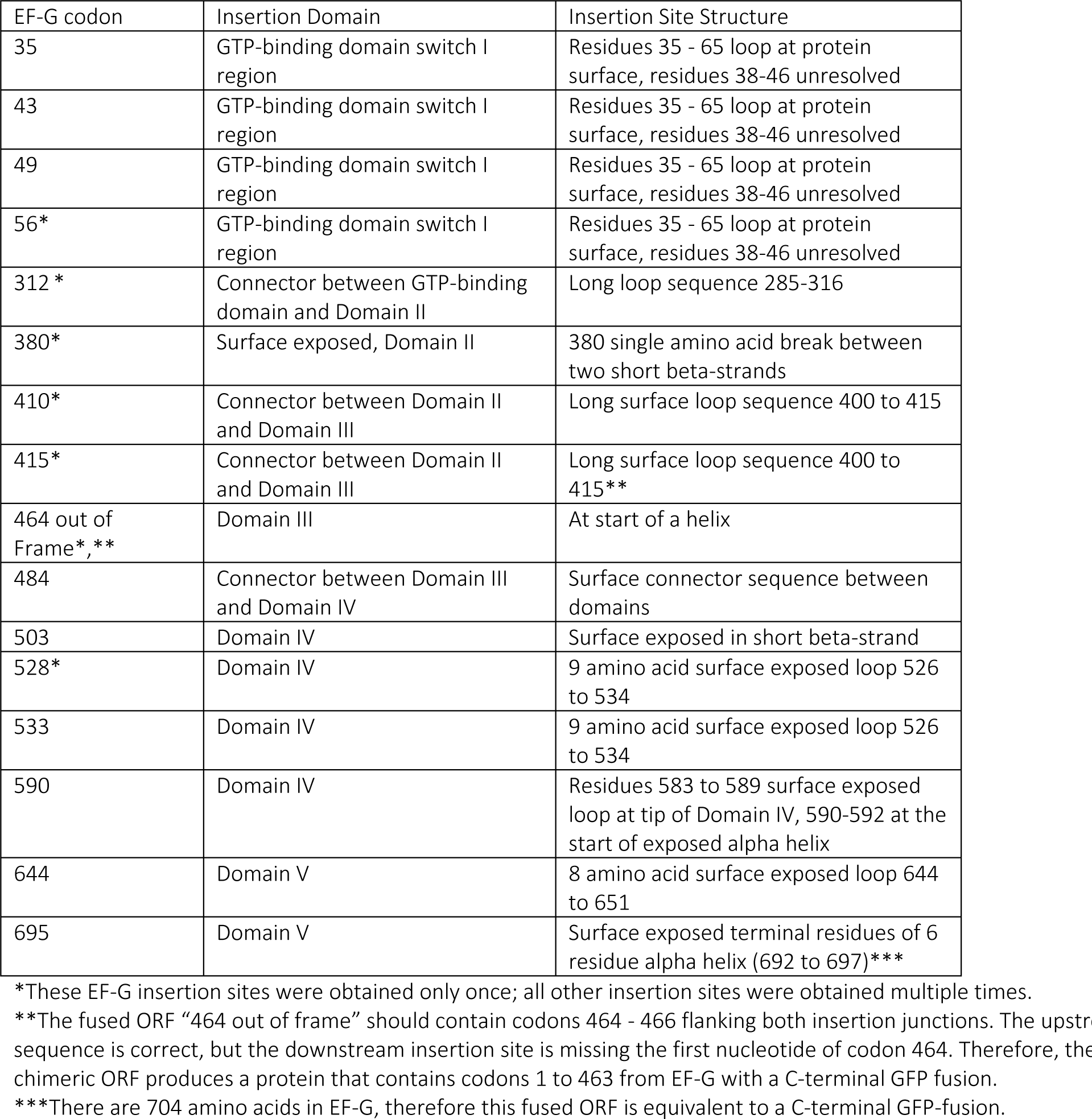
Summary of all insertion points of GFP ORFs into the *fusA* ORF that encodes EF-G. The insertion site structure column provides a description of the structural elements found at the insertion site. The crystal structure of *E. coli* ribosome-bound EF-G was employed for insertion site structure analysis using PDB ID code 4KIY. These results are from a collection 70 candidate plasmids analyzed by DNA sequencing from two different transposition experiments. The transformations of each transposition were standardized to produce 2,000 transformants per plate. Ten plates were produced in each transformation.

**Table S2-2.**
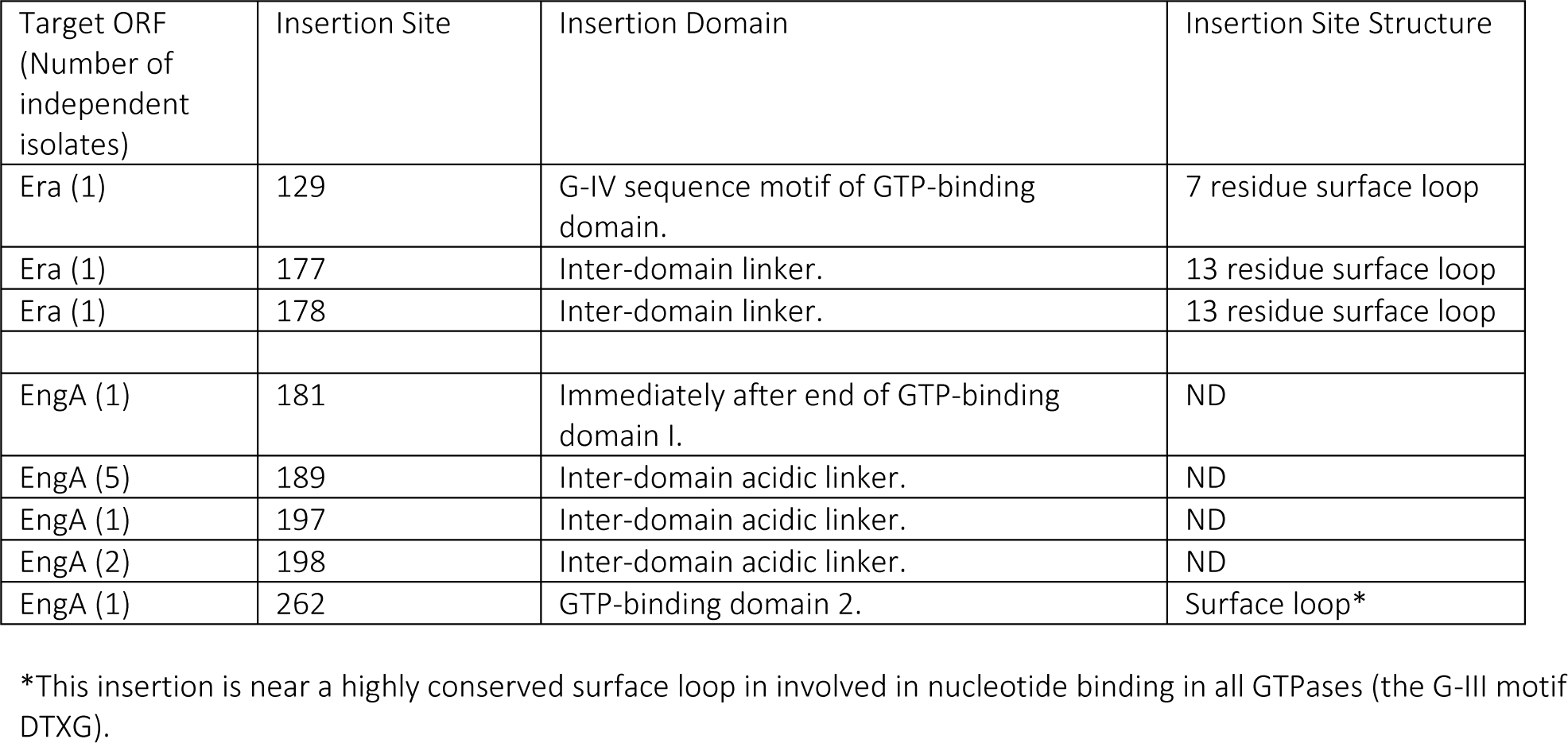
Summary of all insertion points of GFP ORFs into the target ORFs Era and EngA. The insertion site structure column provides a description of the structural elements found at the insertion site. The crystal structure of *E. coli* Era (REF, PDB ID 3IEU) was employed for insertion site structure analysis, but the structure of *E. coli* EngA has not been determined. These data are from one transposition for each target ORF. The transformations of each transposition were standardized to produce 2,000 transformants per plate. Ten plates were produced in each transformation. Ten insertions were identified in the EngA ORF and three in the Era ORF.

**Table S2-3.**
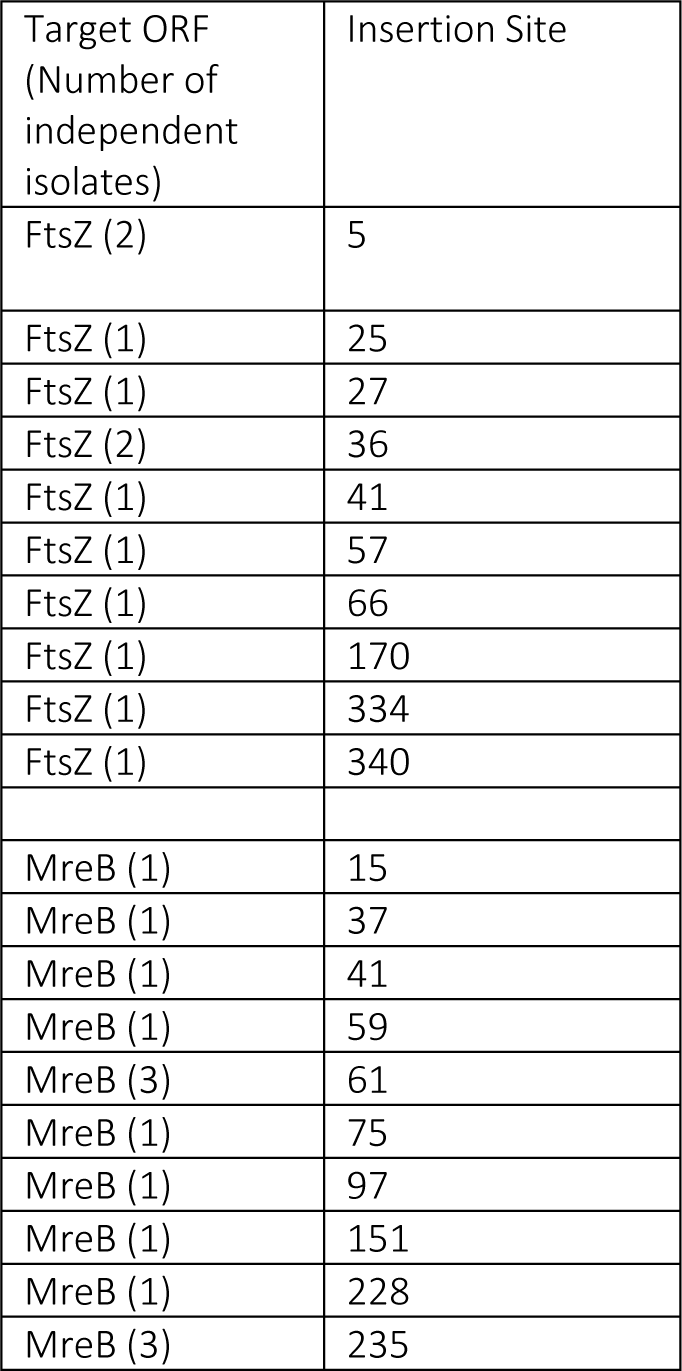
Summary of all insertion points of GFP ORFs into the target ORFs FtsZ and MreB. An exhaustive structural analysis of the insertion sites was not undertaken. To increase the screening capacity two transposition reactions were pooled, and transformants were plated on large petri dishes (150 mm x 15 mm). The transformations were standardized to produce 20,000 transformants per plate. Six plates were produced in each experiment representing a screen of a total of 120,000 colonies per ORF. These data are compiled from using this approach on two separate occasions. Overall, ten insertions into FtsZ and fourteen into MreB were documented.

